# Membrane-embedded polar residues target membrane proteins for degradation by the quality control protease FtsH

**DOI:** 10.1101/2023.12.12.571171

**Authors:** Michal Chai-Danino, Noy Ravensary, Tetiana Onyshchuk, Martin Plöhn, Alon B.D. Barshap, Aseel Bsoul, Hadas Peled-Zehavi, Nir Fluman

## Abstract

Membrane proteins (MPs) navigate challenging biogenesis. Errors in this process are rigorously surveilled by cellular quality control to eliminate faulty MPs. The first critical challenge of this surveillance is the accurate recognition of misfolded proteins. However, how this recognition is achieved for MPs remains poorly defined. Here we reveal the specificity mechanism of FtsH, the major quality control protease clearing faulty MPs in *Escherichia coli*. Analyzing the in vivo degradation of two substrates, we show that lipid-facing polar residues direct substrates to FtsH-mediated degradation. Such polar residues are typically buried in the structural cores of folded MPs, and their exposure to the membrane may thus signify misfolding and flag proteins for degradation. Remarkably, lipid-facing polar residues are sufficient for recognition and can target even folded MPs for degradation. The recognition depends on the FtsH transmembrane domain. Thus, MP misfolding is sensed within the membrane to maintain a healthy membrane proteome.

## Introduction

Every cell faces the challenge of preventing the buildup of misfolded proteins to maintain protein homeostasis. Disruptions to this balance are intimately linked to aging, cancer, and many genetic diseases^1–4^. Protein quality control (QC) systems have been extensively studied for decades. However, the QC of helical membrane proteins (MPs), constituting about a quarter of all proteins^5^, has been relatively neglected until recently^4^. Intriguingly, for several studied MPs, more than 50% of the newly synthesized protein does not fold properly and gets subsequently degraded^6–8^. This suggests that MP QC handles large fluxes. Like their soluble counterparts, misfolded MPs that are not degraded can be toxic to cells^9,10^. On the other hand, excessive degradation of potentially functional MPs is not only wasteful, but can also be detrimental, potentially leading to complete elimination and loss of function of the protein^11,12^. Hence, maintaining the right balance between negligence and over-reactivity is essential. Therefore, a critical step in any QC process is accurately recognizing aberrant misfolded proteins to enable their specific degradation while sparing folded proteins. Despite the critical role of MP QC, our understanding of the molecular mechanisms that mediate misfolded MP recognition is still poor (for more on this, see Discussion).

The FtsH family of proteases serves as the primary MP QC machinery in bacteria, mitochondria, and chloroplasts^13–17^. These well-studied proteins form hexamers composed of two transmembrane helices (TMs) per subunit, and a large soluble domain localized to the cytosol in bacteria^16,18^. The soluble domain is structurally resolved and harbors an AAA+ ATPase that can pull proteins from the membrane and processively feed them into a proteolytic chamber within the hexamer (Fig. 1a)^19–22^. By contrast, The structure of the membrane domain is poorly resolved, and its function beyond serving as a membrane anchor remains unclear^20,23–26^.

**Fig. 1.**
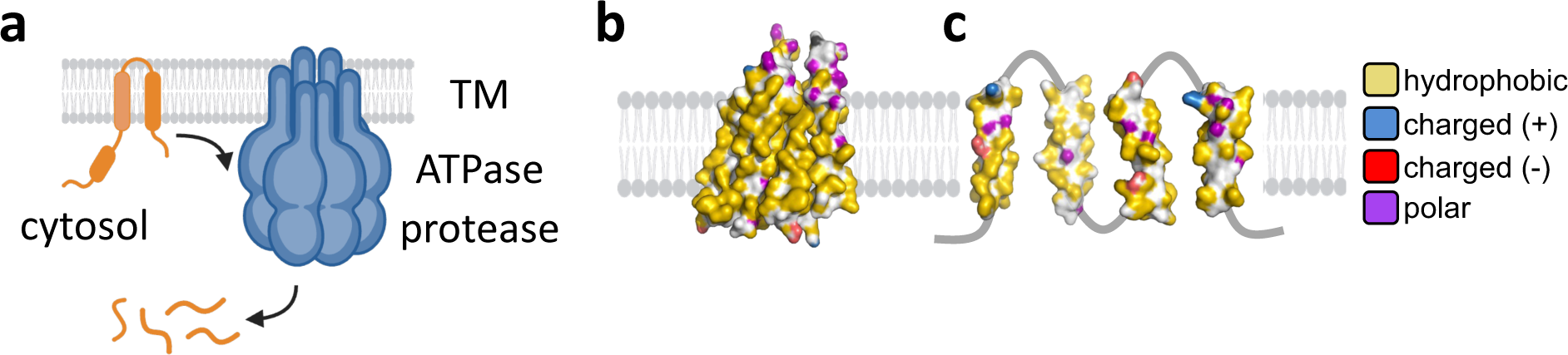
Structural and cellular context of FtsH-mediated MP degradation. **(a)** Illustration of MP degradation by hexameric FtsH. The substrate is pulled from the membrane by the AAA+ ATPase domain and fed into a proteolytic chamber for processive degradation. **(b,c)** Hydrophobic, polar, and charged groups in the predicted MdtJ structure. Hydrophobic carbons that do not lie next to electronegative atoms are colored yellow, polar groups in Ser, Thr, Asn, Gln, and His are colored purple. Charged groups in Asp, Glu, Lys, Arg are colored red or blue, depending on the charge. **(b)** AlphaFold prediction of MdtJ structure, showing the side that faces the lipids in the folded dimer. The lipid-facing surface is hydrophobic. **(c)** Membrane-embedded polar residues in the individual TMs of MdtJ, representing a hypothetical fully unpacked unfolded state. Many polar residues may potentially face the lipid.

Despite decades of study, the mechanism by which FtsH proteases discriminate misfolded from folded MPs remains unknown. Early studies suggested that the *E. coli* FtsH does not possess an unfoldase activity^27,28^, implying that folded proteins would not be degraded. However, these studies were carried out with soluble substrates of FtsH, and later studies with MP substrates revealed that FtsH can, in fact, unfold MPs^29^. Beyond the folding state of the substrate, various substrate sequence attributes were shown to play a role in substrate recognition. These include degrons like ssrA and others^30–32^, as well as long polypeptide stretches that extend out of the membrane and are required to initiate pulling by the ATPase^33,34^. However, these attributes are insufficient in explaining the recognition of misfolded MPs while sparing folded ones. Here we bridge this knowledge gap for the *E. coli* FtsH.

FtsH is a generalist protease, capable of recognizing many misfolded MPs^16,35^. We therefore reasoned that it should recognize some general feature(s) common to misfolding in the membrane. Little knowledge is available on the behavior and structure of misfolded MPs. However, several studies suggest that their transmembrane helices may partially unpack from one another when they are not folded^36–38^. While the extent of unpacking is unclear, this suggests that surfaces and residues that face the core of the structure in the folded protein might be exposed to the lipid upon misfolding. A critical type of residue showing differential exposure to the core versus surface are polar residues. These residues, which are often charged, are present in the transmembrane part of many membrane proteins, often facilitating crucial roles such as ion binding in transporters and channels. However, folded MPs usually bury these residues in the core of the structure, away from lipids, while the lipid-exposed surface remains very hydrophobic (Fig. 1b,c). We hypothesized that the TM unpacking caused by misfolding may expose polar residues to lipids, potentially serving as a signal to sense misfolding by MP QC. Indeed, emerging evidence shows that such polar residues are sensed by MP QC systems in various membranes^39–43^. Here we address the relevance of this hypothesis to FtsH by systematically investigating the *in-vivo* degradation of two substrates. Our results show that membrane-embedded polar residues are essential and sufficient for recognition, and can remarkably target even a folded protein for degradation. We further find that the transmembrane domain of FtsH is crucial for the recognition of polar residues. Our findings provide a clearer picture of the precision and selectivity inherent to cellular quality control.

## Results

### Polar residues are essential for the degradation of orphan MdtJ by FtsH

To investigate how FtsH recognizes misfolded MPs, we initially focused on the dimeric family of Small Multidrug Transporter (SMR). These proteins provide an ideal model, as they remain unfolded until they dimerize, allowing us to control their folding status^36^. In the monomeric unfolded state, their TMs are at least partially unpacked^36^. In addition, FtsH was shown to degrade orphan monomers of an SMR family member^44,45^. To manipulate the dimerization and folding of an SMR protein in a cellular context, we turned to heterodimeric family members, allowing the expression of one subunit without its partner^46^. Indeed, when we expressed the *Lawsonia intracellularis* MdtJ (Uniprot accession, Q1MPU8) as an orphan GFP-MdtJ fusion, it was degraded in an FtsH-dependent manner, with a half-life of about 40 min (Fig. 2a,b,e,f). Co-expression of its cognate dimerization partner, MdtI (Uniprot accession Q1MPU9), abolished the degradation (Fig. 2a,b). Thus, FtsH degrades only orphan and aberrant MdtJ, consistent with QC-targeted selectivity.

**Fig. 2.**
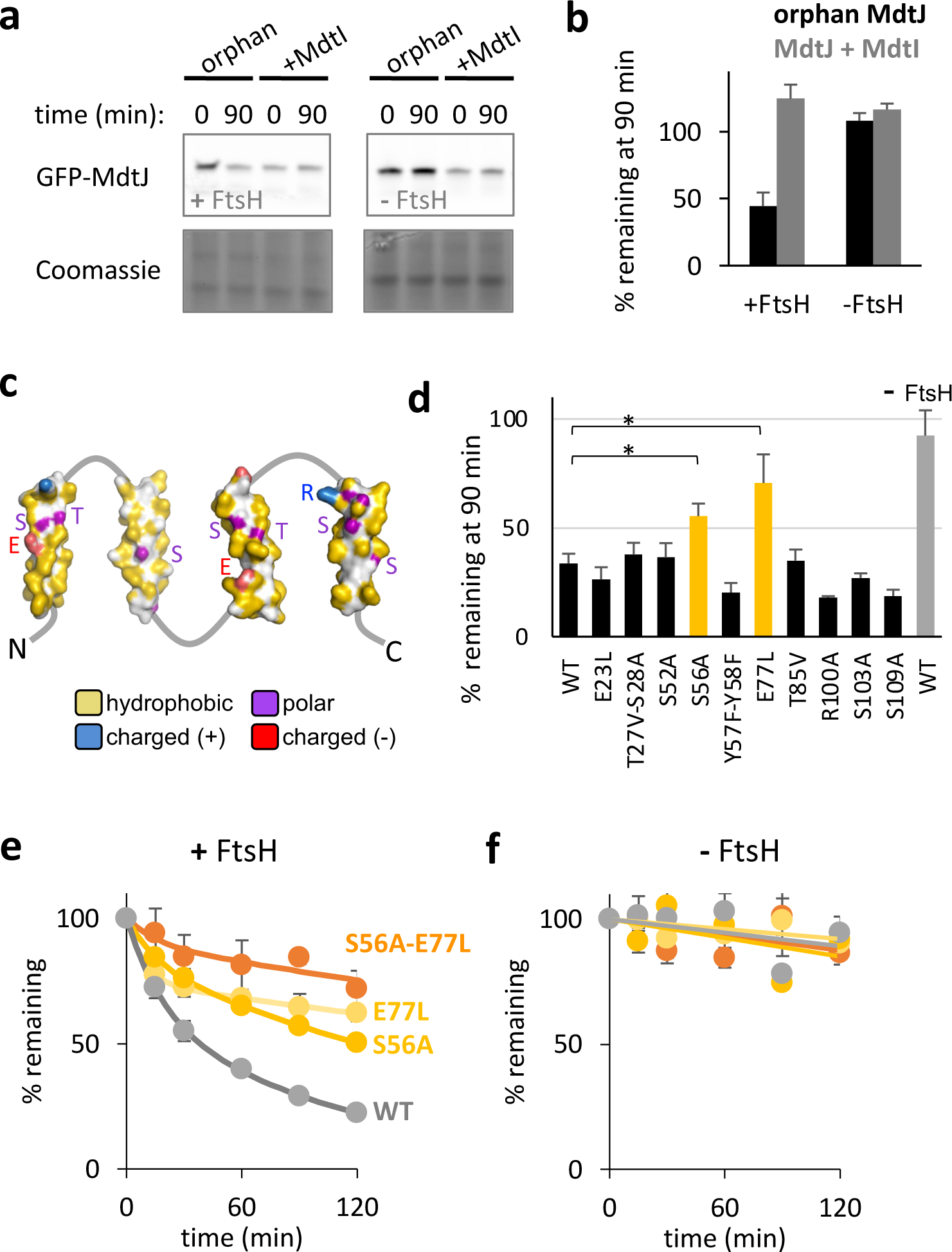
Polar residues are essential for the degradation of orphan MdtJ by FtsH. **(a)** FtsH mediates degradation of orphan GFP-MdtJ. MdtJ was expressed as an orphan protein or with its cognate partner MdtI. At time 0, spectinomycin was added to stop translation and the degradation was assessed after 90 min. In the FtsH knockout strain there is no evident degradation, while co-expressed of FtsH conferred degradation of orphan MdtJ. Coomassie staining of the gels is shown as a loading control. **(b)** Quantification of the experiment shown in (a). Bars show averages of ± SEM of three biological replicates. **(c)** Membrane-embedded polar residues in the individual TMs of MdtJ, which may potentially be recognized by FtsH. **(d)** Effect of substituting polar residues of MdtJ with similarly sized apolar amino acids on degradation, as assessed by the amount of orphan GFP-MdtJ remaining after 90 minutes. Mutations to S56 and E77 perturb the degradation. Degradation of wild-type (WT) GFP-MdtJ in the presence or absence of FtsH are shown as controls. The quantitative analysis is identical to Fig. 2b, from three biological replicates. * indicates p < 0.005 (t-test). **(e)** Degradation time course of GFP-MdtJ mutants in the presence of FtsH, showing that the mutations E77L or S56A slow down the degradation, and their combination almost completely abolishes degradation. Shown are averages ± SEM, as quantified from three biological replicates similar to Fig. 2b. Solid line: the data were fit to a double-exponential equation *Y(t)* = *A_1_*⋅*e*^−*k*1⋅^*^t^* + *A*_2_⋅*e*^−*k*2⋅^*^t^*, where *A*_1,2_ are protein fractions and *k*_1,2_ are rate constants. **(f)** Same as (e), but in the absence of FtsH.

We hypothesized that polar residues within the transmembrane regions of MdtJ may target the misfolded protein for degradation. MdtJ has no solved structure, yet its high-confidence Alphafold prediction is consistent with the solved structures of dimers from the SMR family^46^. According to the predicted structure, the protein has 4 TMs and exposes a hydrophobic surface to the membrane (Fig. 1b). In the misfolded state, these helices may fully or partially unpack from each other, potentially exposing several polar residues to the membrane (Fig. 2c). We mutated each of its transmembrane polar residues, alone or in adjacent pairs, and substituted them by hydrophobic residues of similar size. Most of these residues were dispensable for FtsH-mediated degradation. However, two substitutions, S56A and E77L, significantly slowed down the degradation when mutated alone (Fig. 2d), and almost completely abolished the degradation when mutated together (Fig. 2e,f). These results are consistent with the notion that the TMs of MdtJ at least partially unpack in the unfolded monomeric state, since S56 is predicted to be buried in the core of the folded monomer and would not be accessible for recognition. Interestingly, while S56 and E77 were crucial, other, equally polar residues were dispensable for degradation. This suggests that the positional or structural context of the polar residues plays a role in their recognition, which deserves further study. Nevertheless, the role of S56 and E77 implies that membrane-embedded polar residues are essential for MdtJ recognition by FtsH.

### Lipid-facing polar residues can target a folded protein for degradation by FtsH

Compared to its folded dimeric form, orphan MdtJ is more dynamic^36^ and likely exhibits several structural differences that might distinguish it from folded MdtJ. Therefore, it is plausible that in addition to polar residues, other properties of the misfolded substrate are recognized simultaneously and required for degradation. To examine if polar residues are sufficient on their own, we engineered a *folded* membrane protein to atypically expose a polar residue to the lipid bilayer.

We utilized the *Yersinia fredrixinii* ASBT, a bacterial homolog of the human apical sodium bile acid transporter, whose structure confirms a hydrophobic surface exposed to the lipids^47^. To avoid protein misfolding due to problems in TM insertion into the membrane, we focused on the most hydrophobic ASBT TM3, which is likely robustly inserted. We introduced a mutation in the middle of this TM, at the lipid-facing residue L73, replacing it with a hydrophilic glutamine (Fig. 3a).

**Fig. 3.**
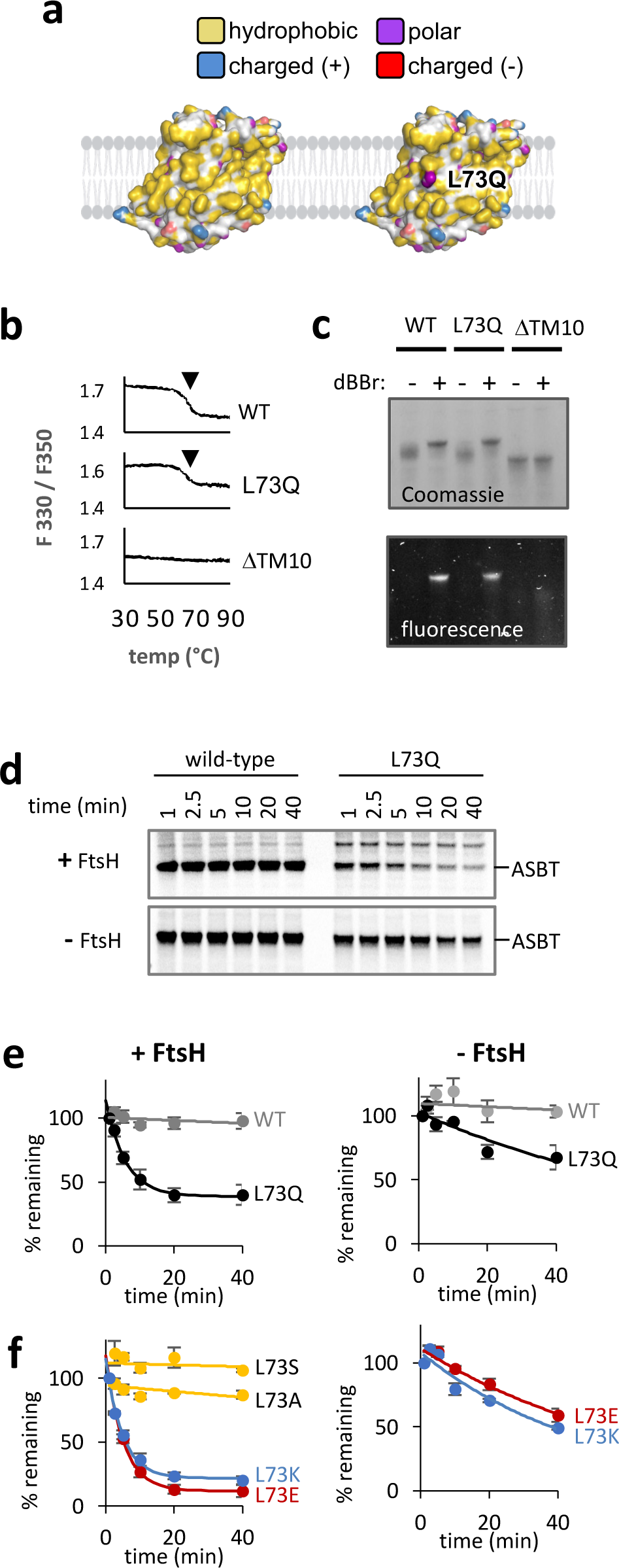
A lipid-exposed polar residue directs folded ASBT for degradation. **(a)** The hydrophobic surface presented to the membrane by ASBT, and the hydrophilic group of glutamine pointing to the membrane in the L73Q mutant. Color coding is as in Fig. 1b,c. **(b)** Thermal denaturation suggests that ASBT-L73Q is stably folded. Purified ASBT and its L73Q and Λ1TM10 mutant were gradually heated, and denaturation was followed by the ratio of fluorescence at 330 and 350 nm. The downward arrow indicates the unfolding transition. **(c)** Crosslinking of ASBT L92C-V219C by dBBr suggests that ASBT-L73Q is folded. Following crosslinking in native *E. coli* membranes, the proteins were purified, run on SDS-PAGE, and detected by Coomassie and in-gel fluorescence. Crosslinking is shown by a shift in gel migration and the appearance of a fluorescent band. **(d)** Following degradation of ASBT and its L73Q mutant by radioactive pulse-chase followed by autoradiography. L73Q is degraded in an FtsH-dependent manner. **(e,f)** Degradation of ASBT and its mutants in the presence and absence of FtsH. Shown are means ± SEM, quantified from three pulse-chase biological repeats (as in d). Solid lines depict nonlinear regression fits to an exponential decay equation where a fraction of the protein remains stable over time. **(e)** FtsH degrades ASBT-L73Q. **(f)** FtsH degrades the charged L73E and L73K mutants, but not the less polar L73A and L73S mutants.

We first confirmed that ASBT-L73Q is properly folded. Wild-type ASBT and its L73Q mutant were purified from FtsH-deleted cells and their thermal denaturation was measured by differential scanning fluorimetry. The mutant and wild-type displayed an unfolding transition at ∼65°C, suggesting similar thermal stabilities (Fig. 3b). As a negative control, we examined an ASBT mutant lacking the centrally-located TM10, which is unlikely to stably fold without the many tertiary interactions provided by this TM (Fig S1a). Indeed, the ΔTM10 mutant showed no indication of thermal unfolding (Fig. 3b). Thus, ASBT-L73Q appears to be stably folded when purified in detergent.

Having established the thermal stability of ASBT-L73Q in detergent, we further probed its folding in native membranes using a crosslinking assay. The assay tests the structural proximity of two residues that would only be within crosslinking distance if the protein is folded. To this end, cysteines were introduced into otherwise cysteine-less ASBT at two positions, L92C and V219C, which lie close in the three-dimensional structure of folded ASBT (Fig. S1b). Several observations confirm that the two cysteines are effectively crosslinked by dibromobimane (dBBr) (Fig. S1c). First, we observed a shift in gel migration of the intramolecularly crosslinked protein (Fig. 3c, lane 2). Secondly, dBBr becomes fluorescent once it reacts with two cysteines simultaneously^48^, and indeed the ASBT-dBBr adduct is highly fluorescent (Fig. 3c, bottom). Thirdly, neither gel-shift nor increased fluorescence was observed in ASBT mutants harboring only a single Cys at position L92C or V219C, confirming that both cysteines crosslink together (Fig. S1d). ASBT-L73Q was fully crosslinked in native membranes, while the misfolding caused by truncation of TM10 abolished crosslinking (Fig. 3c). Considering the location of the crosslinking sites (Fig. S1b,c), these results confirm that the tertiary structure of ASBT-L73Q is maintained in the native membrane. While we cannot exclude a local perturbation to the structure, our results suggest that the L73Q mutant retains its overall structure and stability.

The folded status of ASBT-L73Q suggests that it should be spared by degradative QC. Remarkably, however, this mutant gets rapidly recognized and degraded when expressed in *E. coli*, as if it were an aberrant protein (Fig. 3d,e). Interestingly, nearly 40% of ASBT-L73Q escapes degradation by an unknown mechanism that deserves further research. Deletion of the FtsH gene significantly decreased the degradation rate, but did not abolish it (Fig. 3d bottom, Fig. 3e right panel). Thus, while FtsH appears to be the major protease degrading ASBT-L73Q, other *E. coli* proteases can also recognize membrane-facing polar residues. Additional substitutions in position 73 showed that the polarity of the residue correlated with degradation (Fig. 3f). Relatively apolar substitutions in this position did not lead to degradation, while charged residues, whether negative or positive, elicited an even faster and more complete degradation than the uncharged glutamine. Like the L73Q mutant, the absence of FtsH did not abolish the degradation of the charged mutants completely, suggesting that an additional redundant protease exists (Fig. 3f). Collectively, our findings demonstrate that a single lipid-facing polar residue is sufficient for targeting even a folded MP for degradation, highlighting the sensitivity of the QC system to polar residues.

### Lipid-facing polar residues can trigger the degradation of an assembled dimer

We further tested if membrane-facing polar residues are sufficient on their own to trigger degradation. To this end, we coexpressed GFP-MdtJ together with its partner MdtI to assemble the folded complex. While the lipid-facing surface of folded MdtJ is hydrophobic, we hypothesized that introducing a polar residue facing the lipid might trigger its degradation in the dimer context. Glutamines were introduced at two positions, L55Q and L79Q, lying close in sequence to S56 and E77 that mediate orphan MdtJ degradation. However, unlike these endogenous polar residues, the engineered glutamines are predicted to face the lipids (Fig. 4a). In the absence of FtsH, these mutants had stable expression, with or without MdtI (Fig. 4b). In the presence of FtsH, the co-expression of MdtI stabilized both mutants compared to their orphan counterparts, suggesting that the mutations did not substantially perturb the heterodimer association (Fig. 4c). As discussed below, the intactness of the dimer is further supported by the observation that while expression of orphan MdtJ is toxic to cells, co-expression of MdtI alleviates this toxicity. Remarkably, both mutations triggered the degradation of MdtJ even when co-expressed with MdtI and forming dimers with it (Fig. 4c, gray bars) suggesting that polar residues are sufficient to trigger degradation. Notably, even as orphan proteins, the mutations accelerated the degradation of monomeric MdtJ (Fig. 4c, black bars), suggesting that increasing the polarity of orphan MdtJ enhances degradation. Thus, single glutamines placed at two distinct positions accelerate the degradation of MdtJ by FtsH, both in the context of the orphan and of the assembled MdtJ.

**Fig. 4.**
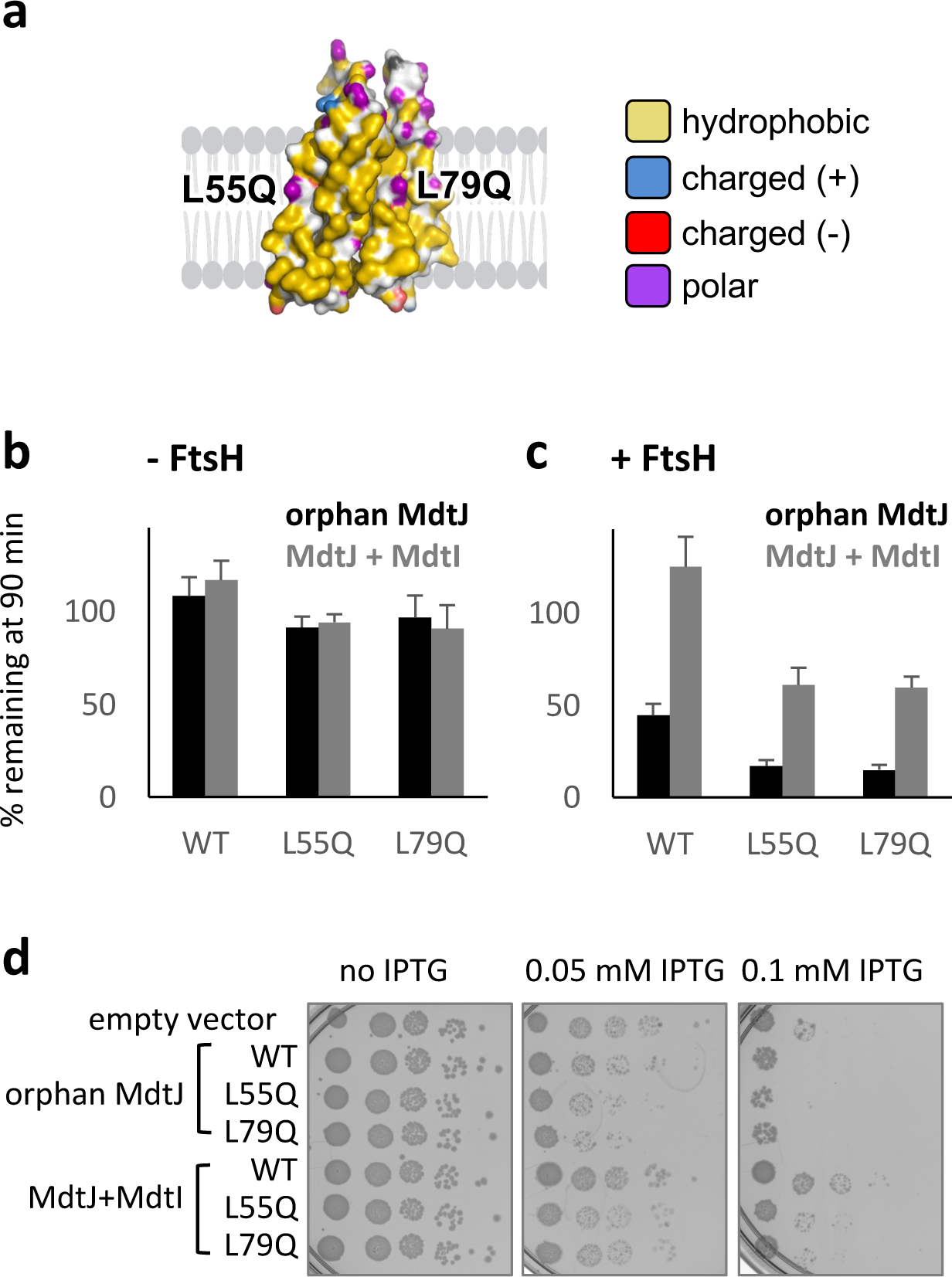
Polar residues can direct assembled MdtJ to degradation. **(a)** The positions of the L55Q and L79Q mutations on the lipid-facing side of MdtJ. The hydrophilic groups of the glutamines are colored purple. Color coding is as in Fig. 1b,c. **(b,c)** Effect of L55Q and L79Q mutations on GFP-MdtJ degradation in the absence (b) and presence (c) of FtsH. Bars show averages of ± SEM of three biological replicates. **(d)** Effect of polar residues of MdtJ on *E. coli* growth. A serial 10-fold dilution of bacterial culture was spotted on LB-agar plates under no induction (no IPTG) or under increasing IPTG-controlled induction. The level of toxicity is inferred by reduced growth compared to cells harboring an empty vector.

Interestingly, the polarity of MdtJ correlated with the toxicity conferred by its expression in *E. coli*. We tested the effect of IPTG-controlled MdtJ expression on *E. coli* growth. Wild-type MdtJ was toxic when induced, whereas co-expression of MdtI alleviated this toxicity (Fig. 4d), consistent with misfolded orphan MPs being harmful to the cell. We postulated that the lipid-facing polar residues, which are sequestered in the assembled dimer, could contribute to the toxicity of the orphan MdtJ. Indeed, when expressed as orphans, both MdtJ polar mutants, L55Q and L79Q displayed toxicity already at lower IPTG induction (Fig. 4d), suggesting that increased polarity exacerbates their toxicity. Moreover, co-expression of MdtI could not fully alleviate the toxicity of both mutants (Fig. 4d), suggesting that the lipid-facing glutamines of the L55Q and L79Q mutants are toxic in the dimer context. Thus, lipid-facing polar residues may be toxic to the cell, explaining why proteins bearing them need to be eliminated by QC.

### The FtsH transmembrane domain participates in MP QC

We next asked how FtsH recognizes substrates bearing membrane-facing polar residues. Since the recognition event likely takes place in the membrane, we reasoned that the transmembrane domain of FtsH might be involved. FtsH possesses two TMs, and the membrane-anchoring provided by the TMs was shown to facilitate hexamerization^26^. However, beyond the roles in anchoring and hexamerization, studies of various FtsH homologs yielded conflicting results on the importance of its TMs. In *S. cerevisiae,* the TMs appear important for the degradation of some integral MPs^20,25^. By contrast, the TMs of the *E. coli* FtsH appear dispensable, as swapping them with unrelated TMs from the LacY protein retains the degradation of orphan SecY^26^. We speculated that FtsH, as a major MP QC factor, might have multiple modes of recognition for different MP substrates. Therefore, we set out to investigate the role of the TMs in the recognition of membrane-facing polar residues.

We first repeated the approach of Akiyama and Ito, swapping the N-terminal, TM-containing part of FtsH, with TMs 1 and 2 of LacY^26^ (Fig. 5 a,b). Two LacY-FtsH chimeras were generated, one with the wild-type LacY TMs and another more membrane-inert version, called hydrophobic LacY-FtsH, where the polar residues in LacY TMs were replaced by apolar ones (Fig. 5b). We utilized FtsH self-processing to examine if these chimeric versions retain the hexameric structure needed for protease activity. Briefly, FtsH self-processing refers to the enzyme’s ability to cleave its own C-terminus, a process that does not affect its protease activity^49^. Indeed, when FtsH-3xHA was expressed, the HA-tag was readily trimmed, regardless of the swapping of N-terminal TMs (Fig. 5c), indicating that the LacY-FtsH chimeras remain hexameric active proteases. We further confirmed that both LacY chimeras retained degradation activity towards a soluble substrate, the phage Lambda regulatory protein CII^50^ (Fig. S2), suggesting that the general activity of FtsH remains intact. Nevertheless, neither LacY-FtsH versions had activity towards orphan MdtJ, suggesting that the transmembrane domain is essential for degradation of MPs bearing lipid-facing polar residue (Fig. 5c,e).

**Fig. 5.**
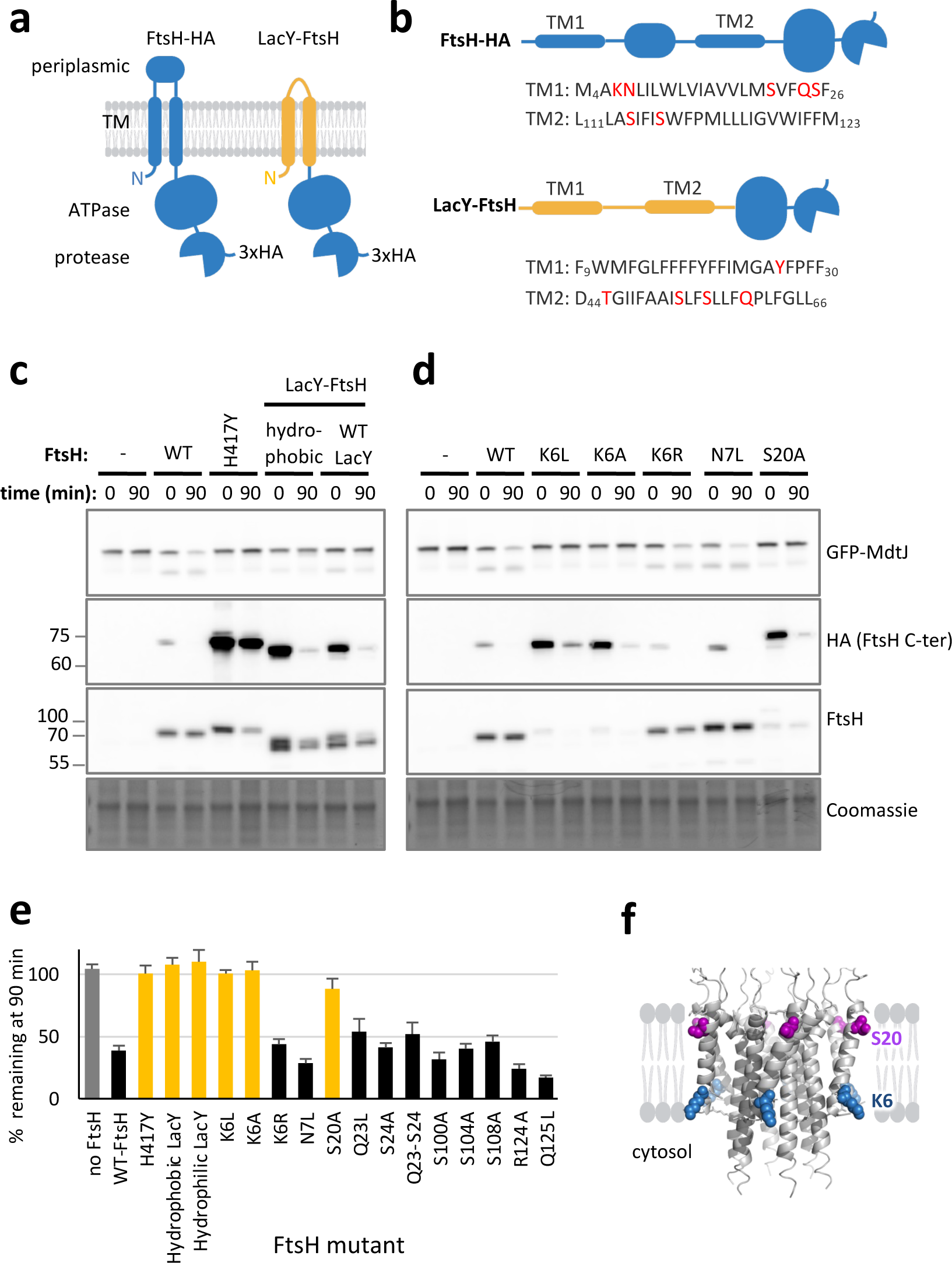
The FtsH transmembrane domain facilitates MP QC. **(a)** Topological representation of FtsH and LacY-FtsH, both with a C-terminal HA tag. **(b)** Sequences of TMs 1 and 2 of FtsH and LacY. The highlighted residues were mutated when indicated. FtsH residue numbering is as in Wang et al (1998), adding 3 residues to the N-terminus of FtsH. The mutations to LacY TMs, generating ‘hydrophobic LacY’ were Y26V, T45V, S53A, S56A, and Q60A. **(c,d)** Activity of FtsH variants in degradation of MdtJ and of the FtsH C-terminal HA tag. The degradation experiment was done similarly to Fig. 2a, with additional detection of FtsH and its C-terminus by FtsH and HA antibodies, respectively. H417Y is a catalytically inactive FtsH mutant. Coomassie staining is shown as a loading control. (c) LacY-FtsH chimeras are active proteases, as observed by self-processing of the HA-tag, but cannot degrade MdtJ. (d) Mutations in K6 and S20 cause pleiotropic effects in FtsH, resulting in reduced expression, FtsH degradation, and loss of activity. **(e)** Quantification of MdtJ degradation by various FtsH variants, with apolar mutations to polar TM residues. Most polar residues are dispensable for MdtJ degradation, except for swapping the TMs of FtsH with LacY TMs 1+2, and mutations to K6 and S20. Shown are averages ± SEM from at least three biological replicates. Representative gels are shown in panel c,d and Fig. S4. **(f)** Location of S20 and K6 in the AlphaFold model of hexameric FtsH, showing that they are predicted to face the membrane, near the interface.

We then hypothesized that FtsH might directly recognize membrane-embedded polar residues through its transmembrane domain. The FtsH transmembrane domain is poorly resolved in the current structures^22^, suggesting considerable dynamics of this part of the protein. We generated an AlphaFold model of the hexameric N-terminal part of FtsH. We obtained relatively low confidence in most of the transmembrane parts (pLDDT between 50 and 70, Fig. S3), supporting the notion that the TMs are dynamic and indicating that the model should be taken with caution. Nevertheless, the prediction shows good agreement with the predicted TM boundaries of FtsH and a part of TM2 that was resolved in a recent structure^22^, suggesting that it well captures the hexameric core and topology.

We then replaced all polar residues within the FtsH TMs, individually or in adjacent pairs, with similarly sized apolar ones (residue numbering was as in Wang et al.^51^, adding 3 residues to the N-terminus). Most of these mutants retained full activity, both towards MdtJ and self-processing (Fig. 5d,e, Fig. S4), suggesting that the polarity of the residues plays no role. However, two mutations, S20A and K6L, perturbed FtsH. The positive charge of K6 appeared important, as replacing it with a positively charged arginine preserved activity, while a neutral alanine or leucine abolished it (Fig. 5d,e). Notably, the inactive S20 and K6 mutants displayed reduced cellular stability, as indicated by degradation of the full-length FtsH (Fig. 5d, FtsH panel). The S20A FtsH mutant also unexpectedly migrated more slowly in SDS-PAGE, suggesting an unexplained apparent mass shift (Fig. 5d, HA panel). These mutants, therefore, have pleiotropic effects on FtsH function, likely reflecting a general role in the protein, leaving it impossible to discern whether K6 and S20 play a specific role in substrate recognition. Taken together, while the FtsH transmembrane domain appears essential for the recognition of membrane-embedded polar residues, most polar residues appear dispensable for this recognition, at least when mutated individually.

## Discussion

Various mechanisms have evolved to recognize and degrade misfolded MPs. Misfolded MPs can sometimes be recognized by soluble QC factors, if the protein exposes a recognizable misfolded domain outside the membrane^52,53^. However, many MPs, especially multispanning ones, form folded domains or assemble inside the membrane. The misfolding of such membrane-embedded domains likely requires recognition from within the membrane.

One group of factors acting within the membrane are intramembrane proteases, capable of cleaving peptide bonds within transmembrane helices. These proteases are often involved in signaling^54^, yet several were shown to play a QC role by cleaving misfolded proteins^55–59^. The catalytic mechanism of intra-membrane proteases employs a principle that offers selectivity towards misfolded proteins. To cleave a peptide bond, the substrate’s transmembrane helix must transiently unwind to allow the active site to access the polypeptide backbone^57,60–64^. These enzymes, therefore, typically cleave at points where the helix is unstable. Proper folding and assembly of MPs would stabilize the helices against unwinding, allowing the proteins to evade degradation^56,57,63^. An analogous mechanism has recently been shown for Signal Peptidase, which cleaves at a membrane-juxtaposed position^65^. Importantly, the specificity of the cleavage sites means that most TMs cannot be cleaved by these enzymes. Therefore, in the case of multispanning proteins, these proteases leave large membrane-embedded fragments that require further degradation^56,58,63,65^.

The last and most prominent route for MP degradation is by machineries that identify misfolding within the membrane, but lead to complete cleavage of the substrate into peptides. In the eukaryotic endomembrane system, the pivotal step marking a protein for degradation is ubiquitination, which then targets the MP for either extraction from the membrane and degradation by cytosolic proteasomes^66^, or degradation by the autophagy-lysosome pathway^67^. The factors responsible for selectivity in these cases are E3 ubiquitin ligases and their adaptors. However, how these factors specifically target misfolded MPs is poorly understood^66^. For several of these systems, the TMs of either the substrate or the E3 ligase or adaptor were shown to be crucial, suggesting that the recognition takes place within the membrane^39,40,68–72^.

The analogous degradative systems in bacteria and endosymbiotic organelles are FtsH-family proteases. FtsH functions as an all-in-one machine, capable of recognizing a protein, extracting it from the membrane, and processively degrading it into peptides^16^. Notably, while other proteases in *E. coli* are capable of degrading misfolded MPs^35,57,73^, FtsH is assumed to be the major MP QC protease, and is the only processive AAA+ protease anchored to the membrane^16,35,74^. Despite the essentiality of these proteases, the mechanism behind their selectivity towards misfolded MPs remained mysterious.

Here, we revealed that lipid-facing polar residues are recognized by FtsH. This provides an explanation for the specificity towards misfolded MPs, since folded and assembled MPs typically bury membrane-embedded polar residues in their core, away from the lipids. An analogous but mirror-imaged principle guides the QC of soluble proteins, where the aberrant exposure of core hydrophobic residues attracts QC^75^. Moreover, our findings suggest that MPs exposing polar residues to the lipids are toxic to the cell, explaining the need for their elimination by QC. Polar residues are not only essential for recognition, but they are sufficient on their own, allowing us to engineer MPs that get degraded despite being folded. A single glutamine decorating the surface of a folded MP is sufficient to cause its degradation at rates comparable to or even faster than a genuine misfolded protein. This observation showcases the extreme sensitivity of the QC system to lipid-facing polar residues. Interestingly, the finding that many endogenous polar residues of MdtJ were dispensable for degradation suggests that the position of the polar residue plays a role in the recognition. While the mechanistic explanation for this positional effect is unclear, it is plausible that the different positions display variable levels of lipid-exposure in the dynamic misfolded state^76^. Beyond FtsH, polar residues are emerging as a strong recognition determinant for QC in the membrane. Recent evidence suggests that this principle drives recognition by ER membrane chaperones^41–43,77^, an intramembrane protease^56^, and Golgi-localized E3 ligases^39,40^. Interestingly, we find that the degradation of ASBT mutants is not fully abolished in the absence of FtsH, suggesting the existence of additional mechanisms that recognize such polar residues in *E coli*.

How does FtsH recognize polar residues in the membrane? Our findings indicate that the FtsH TMs play a role in this recognition, but the fine mechanistic details remain unclear. One possibility is a direct interaction between substrates and FtsH, potentially via membrane-embedded polar residues of FtsH. While we did not find any polar residue whose mutation affected degradation, the possibility of direct interaction is still plausible. Firstly, it is possible that several FtsH polar residues can contribute to the interaction and redundantly compensate for the mutation of single residues. Moreover, the residues that are structurally most likely to interact with the substrate, K6 and S20, which face the membrane near its interface (Fig. 5f), could not be studied because their mutants were degraded in *E. coli*. This destabilization is somewhat surprising since increasing the hydrophobicity of lipid-facing residues is expected to stabilize the protein. The results suggest that these residues play an important functional role in FtsH that deserves further investigation. The FtsH lipid-facing polar residues may play QC roles beyond substrate recognition, as they can potentially cause a membrane perturbation around the protease. Such a perturbation may either facilitate MP extraction^78^, or aid colocalizing FtsH to areas of thinned bilayers, where misfolded MPs might reside. An alternative possibility is that FtsH does not recognize polar residues directly, but requires an adaptor. Indeed, FtsH was shown to collaborate with a number of adaptors^31,58,79,80^. However, FtsH has multiple substrates, both membrane and soluble proteins ^15,22,31,32^, requiring different recognition modes, and some substrates can be recognized directly by FtsH^29,32,81^.

To conclude, our study clarifies how FtsH distinguishes folded from misfolded MPs and opens the door for further investigation into how lipid-facing polar residues are engaged by this protease. This recognition mechanism, focusing on a feature likely common to misfolded MPs, may endow FtsH and similar quality control factors with the ability to target a diverse array of substrates. The emerging recognition of membrane-embedded polar residues as key determinants in protein quality control marks a significant step forward in decoding the complex language of membrane protein homeostasis.

## Materials and Methods

### Bacterial strains

*E. coli* DH5α was used for plasmid propagation. BL21 (DE3) and the Δ*ftsH* strain AR5090 (DE3)^45^ were used for protein expression. Radioactive pulse-chase degradation assays of ASBT were done with *E. coli* AR5090 (DE3)^45^ or *E. coli* BL21 (DE3). Toxicity assays of MdtJ protein were done using *E. coli* BL21 (DE3) *ΔemrE::kana^R^, ΔmdtJI::Cm^R^(*ref ^45^), in which deletion of the *E. coli emrE* gene and *mdtJI* operon was used to avoid potential interactions between the expressed proteins and chromosomally encoded SMR transporters.

### Plasmids

All mutants were engineered by PCR-based techniques such as QuickChange and Gibson assembly, and their sequences were verified. pZ ASBT, a pET derivative expressing cys-less ASBT, was described in Zhou et al. 2014^47^, it was further engineered to replace the kanamycin resistance gene with ampicillin resistance. pBAD/HisB-sfGFP (TIR^STD^) was a kind gift from Daniel O. Daley^82^. The pET19b (Novagen) expressing MdtJ and MdtI were described in ref^45^. Both MdtJ and ASBT were Cys-less versions.

### Cloning

The FtsH gene was amplified from the MG1655 *E. coli* strain genome. The amplification included a 200 bp region upstream of the gene, serving as the native promoter, and a 110 bp region downstream, acting as the natural terminator. The following primers were utilized for the amplification: Forward primer: AAATTCGCTCCCTGTTTACGAAGGTC. Reverse primer: ACCCTGGGCAAAGAGTTTCATGATG. After amplification, the gene was cloned into the pACYC184 plasmid using the Gibson Assembly method^83^. Additionally, a triple-hemagglutinin (3xHA) tag was appended to the C-terminus of the protein, facilitating subsequent detection.

The MdtJ and MdtI genes were amplified from the pET 19b MdtJI plasmids. Post-amplification, they were cloned into the pBAD/HisB-sfGFP (TIR^STD^) vector utilizing the Gibson Assembly method. The primers used for the cloning process were as follows:

MdtJ Forward Primer: GATTACACATGGCATGGATGAACTCTACAAAATGCAACCCCGTCAATATACTTCAC.

MdtJ Reverse Primer: CAAAACAGCCAAGCTTCGAATTCTTATTAGTTATTTGATGAAATATTTACTG.

pBAD forward primer: TAAGAATTCGAAGCTTGGCTGTTTTGG. pBAD reverse Primer: TTTGTAGAGTTCATCCATGCCATGTG. Similarly, the MdtI gene was extracted from the pET19b MdtJ+MdtI plasmid.

List of plasmids used:

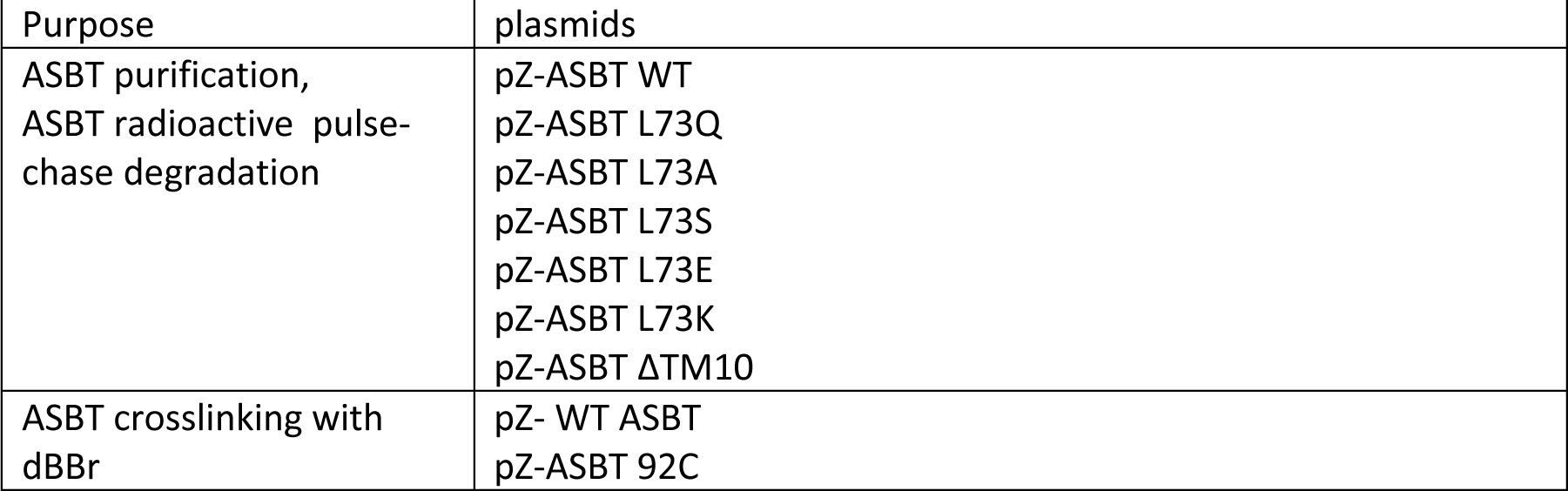
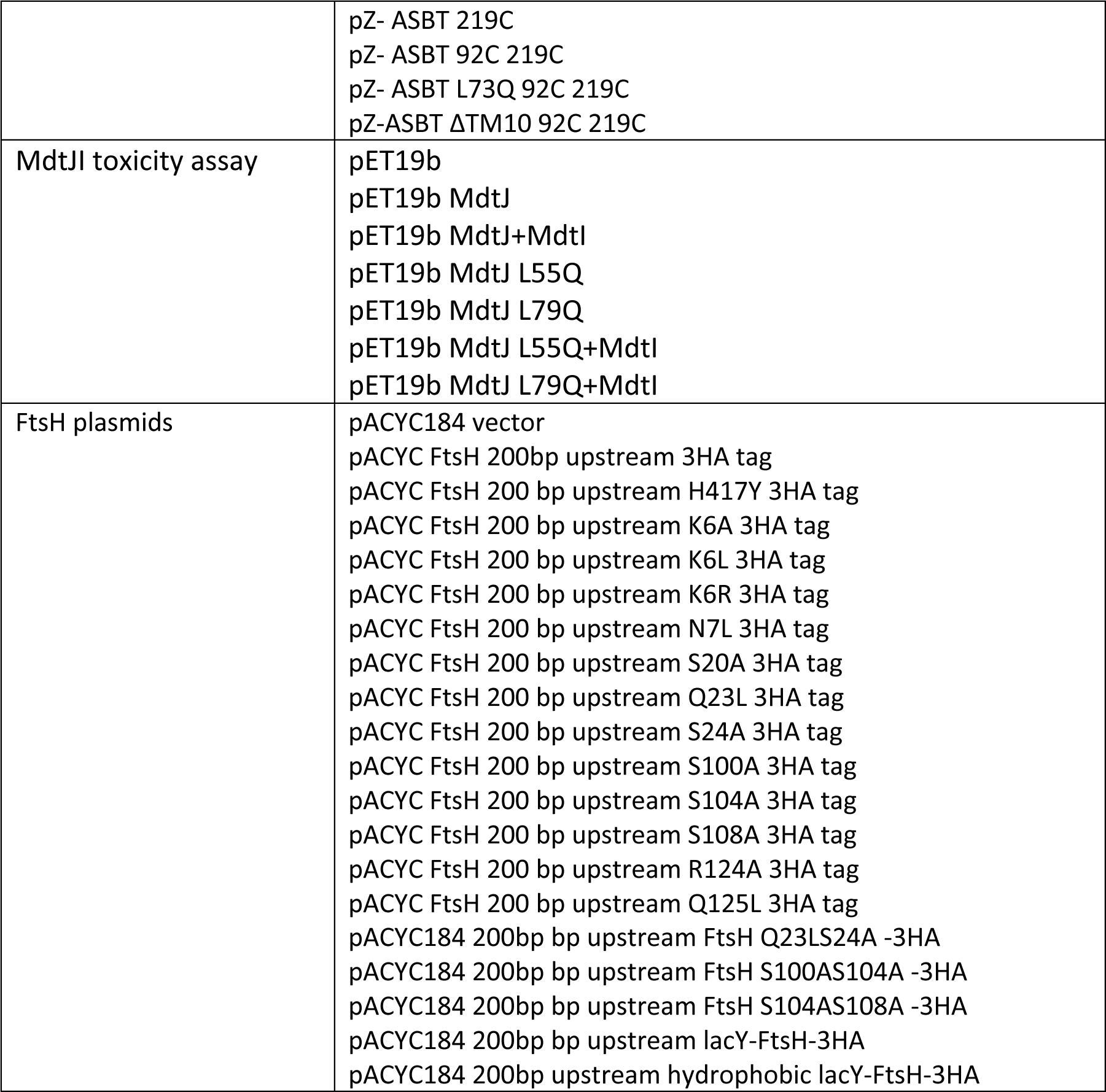

### ASBT degradation by radioactive pulse-chase

A smear of colonies from a fresh transformation of AR5090(DE3) *ΔftsH3::kan, sfhC21/FʹlacI*I^45^ or BL21(DE3) harboring the indicated pZ plasmids was grown overnight in M9 medium supplemented with 1 g/L CSM-Met (MP biomedical, cat 4510712), 100 μg/mL thiamine, 0.4% glycerol, 2mM MgSO_4_, 0.1mM CaCl_2_, 0.5% glucose and 100 μg/mL ampicillin. The culture was then back-diluted to 0.1 OD into the same medium without glucose and grown at 37°C to mid-log phase (OD_600_ of ∼0.5). Cultures were induced with 0.1 mM IPTG for 10 min, followed by 15 minutes of incubation with 0.2 mg/mL rifampicin to halt transcription other than T7 polymerase-dependent transcription. Proteins were labeled with 15 μCi [^35^S]Met for 1 min, then mixed with a high excess (2 mM) of non-radioactive methionine. Samples of 0.3 mL were taken at variable chase times and mixed with 0.75 mL ice-cold PBS to rapidly cool them. Cells were harvested (5 minutes, 10,000*g* at 4°C) and resuspended in 100 μL sample buffer. Protein separation was conducted using 12% polyacrylamide gel electrophoresis. Gels were subsequently dried, visualized by autoradiography, and quantified.

### CII Degradation analysis

The procedure was analogous to the one described for the ASBT mutants. However, the E. coli AR5090 (DE3) *ΔftsH3::kan, sfhC21/FʹlacIq* strain was co-transformed with the pZ-cII plasmid ad the specified pACYC FtsH plasmid. The growth medium was supplemented with 0.5% glucose to reduce CII expression. The appropriate antibiotics were included based on the plasmid resistance marker. Proteins were labeled 15 μCi [^35^S]Met for 1 min and chased for the indicated times.

### MdtJ toxicity and expression assay

A smear of colonies from a fresh transformation of BL21 (DE3) *ΔemrE::kana^R^, ΔmdtJI::Cm^R^(*ref ^45^) harboring the indicated pET19b plasmids, were grown overnight in LB medium supplemented with 100 µg/mL ampicillin. The culture was then back-diluted to 0.1 OD into the M9 medium supplemented with 1 g/L CSM-Met (MP biomedical, cat 4510712), 100 μg/mL thiamine, 0.4% glycerol, 2mM MgSO_4_, 0.1mM CaCl_2_, 100 μg/mL ampicillin) and grown at 37 °C to mid-log phase (OD_600_ of ∼0.5).

For expression analysis by radioactivity, equal amounts of cells (0.5 mL) were divided to 2mL tubes. The tubes were then incubated in a thermoshaker set at 37°C with agitation at 700 RPM. Cultures were induced with 0.1 mM IPTG for 10 min, followed by 15 min incubation with 0.2 mg/mL rifampicin to halt transcription other than T7 polymerase-dependent transcription. Proteins were labeled with 15 μCi [^35^S]Met for 2 min, chased with a high excess (2 mM) of non-radioactive methionine and put on ice. The cultures were harvested (5 min, 10,000*g* at 4°) and the pellet was resuspended in 100 µL of sample buffer. Protein separation was conducted using 12% polyacrylamide gel electrophoresis. Gels were subsequently dried and visualized by autoradiography.

For toxicity assay, five tenfold serial dilutions were prepared from the mid-log cultures, with the highest density having an OD_600_ of 0.1. The serial dilution was spotted (3 µL) on LB agar–ampicillin plates containing indicated amounts of IPTG or glucose. The plates were allowed to grow for one or two nights at 37 °C.

### MdtJ degradation assay

A smear of colonies from a fresh transformation of the FtsH deletion strain AR5090 (DE3) harboring the indicated pACYC and pBAD plasmids, encoding FtsH and GFP-MdtJ, respectively, was grown overnight in LB medium supplemented with 50 µg/mL ampicillin, 17 µg/mL chloramphenicol and 0.5% Glucose. The cultures were then back-diluted to 0.1 OD into the same medium without glucose and were grown at 37 °C to mid-log phase (OD_600_ of ∼0.5). Protein expression was induced by adding 0.1 mM arabinose for 30 minutes. Samples (0.5 mL) were collected and immediately cooled on ice. Cultures were further treated with 200 µg/mL spectinomycin to halt translation and additional sets of samples were collected in the indicated times. Cells were harvested (5 min, 10,000*g* at 4°), washed with 0.6 mL PBS, resuspended with 150 µL lysozyme buffer (150 mM NaCl, 30 mM Tris–HCl pH 8, 10 mM EDTA, 1 mM DTT, 1 mg/mL lysozyme and 1x cOmplete Protease Inhibitor, roche) and freezed at -20°C. Cells were disrupted by thawing at 25 °C for 5 minutes, followed by shaking at 37 °C for 10 minutes. Then, 0.9 mL of DNase solution (15 mM MgSO4, 1 mM DTT and 1 mM PMSF, 1 x Turbo DNase (Jena biosciences)) was added and the samples were allowed to shake at 37 °C for 10 min before transferring to ice. Crude membranes were collected by centrifugation at 4 °C, 20,000*g* for 20 min and the pellet was resuspended in 50 µL of GFP sample buffer (50 mM Tris 8.0, 2% SDS, 10% glycerol, 20 mM DTT, bromophenolblue). Proteins were separated by 12% polyacrylamide gel electrophoresis. Gels were visualized by Typhoon and quantified followed by Coommasie staining. FtsH visualization was done by western blotting with anti-HA antibody (Sigma-Aldrich, H9658) or anti-FtsH antibodies (the National BioResource Project (NBRP)-E. coli, Japan)

### Software

Gels were quantified by the ImageQuant^TM^ software (Cytiva), following the developer’s instructions. Protein structures were visualized using The PyMOL Molecular Graphics System, Version 2.0 Schrödinger, LLC. Hydrophobic surface coloring was using a script generated by Hagemans et al.^84^ Illustrations were Created with BioRender.com.

### ASBT purification and thermal stability assay

ASBT purification was done similarly to Zhou et al.^47^ with slight changes.

Cellular Growth and Harvesting: The indicated plasmids were transformed into the FtsH deletion strain AR5090 (DE3). A single colony was grown in Luria-Bertani (LB) medium supplemented with 0.5% Glucose overnight and then back diluted 100 fold to 1000 mL of the same medium. Cells were grown to an optical density at 600 nm of ∼0.6 at 37 °C. Overexpression of the protein was induced by addition of isopropyl β-D-1-thiogalactopyranoside (IPTG) to a final concentration of 0.5 mM at 20 °C overnight. An equal volume of cells corresponding to 500 OD units was collected and subjected to a washing step with 100 mL of wash buffer (20 mM HEPES, pH 7.5, 150 mM NaCl). Subsequently, the cells were pelleted and resuspended in 50 mL of lysis buffer (20 mM HEPES, pH 7.5, 150 mM NaCl, 10%(v/v) glycerol, 2mM β-mercaptoethnaol)

Membrane preparation: cells were thawed in a water bath, added with Turbo DNase and 1mM PMSF and lysed using emulsiflex 3 times. The lysed cells were centrifuged to discard cell debris. The supernatant was collected and centrifuged to collect membranes (100,000*g*, 60 min at 4°). The pellet containing membranes was resuspended using homogenizer in 8 mL lysis buffer.

Aliquots of 2 mL were snap frozen.

ASBT purification: The supernatant was loaded onto a talon metal affinity column, washed twice with 3 mL wash buffer (20 mM HEPES, pH 7.5, 150 mM NaCl, 10%(v/v) glycerol, 2mM β-mercaptoethnaol, 20 mM imidazole 0.2% DDM) and eluted with elution buffer (20 mM HEPES, pH 7.5, 150 mM NaCl, 10%(v/v) glycerol, 2mM β-mercaptoethnaol, 300 mM imidazole 0.2% DDM).

Protein concentration was measured by nanodrop and calculated by dividing by the extinction coefficient (1.19 M^-1^ cm^-1^ for WT and Glutamine mutant, 0.943 M^-1^ cm^-1^ for ΔTM10). Protein concentrations obtained were 0.2-0.4 mg/mL. Equal amounts of protein were dialyzed twice for at least 1 hour each time in lysis buffer .

The thermal stability was assayed for 2 replicas in a volume of 10 µL each. Changes in the intrinsic fluorescence were measured by nanoDSF (Prometheus NT.48) . Lysis buffer supplemented with 0.2% DDM served as blank.

### Crosslinking by dibromobimane (dBBr)

Cell growth and harvesting: 500 mL of cells were grown, harvested and washed as described in the previous section and resuspended in in 50 mL lysozyme buffer (150 mM NaCl, 30 mM Tris–HCl, pH 8.0, 10 mM EDTA, 1 mg/mL lysozyme, phenylmethanesulfonyl fluoride (PMSF)). The suspension was snap-frozen.

Crude membrane preparation: The frozen cells were thawed in a room-temperature water bath and then moved to 37°C shaker with intermittent shaking for 10 minutes. 180 mL of DNase solution (15 mM MgSO_4_, Turbo DNase (Jena biosciences), 0.5 mM PMSF) was added, and the mixture was further agitated for 15 minutes. After incubation, the cell suspension was immediately transferred to an ice bath for 10 minutes.

Crude membranes were isolated by centrifugation at 20,000*g* for 30 minutes at 4°. The resulting pellet was resuspended in 3.8 mL of ASBT buffer (20 mM HEPES, pH 7.5, 150 mM NaCl, 10% v/v glycerol, 2 mM β-mercaptoethanol) using a homogenizer.The membrane suspension was aliquoted into six separate tubes and rapidly frozen in liquid nitrogen for subsequent analyses.

Membrane washing and crosslinking: One tube for each mutant was thawed in a water bath. The membranes underwent two wash cycles using 1 mL of membrane wash buffer (20 mM HEPES pH 7.5, 150 mM NaCl) to remove any residual reducing agents. Subsequently, membranes were resuspended in 1.1 mL of the same buffer. Aliquots of 500 µL were divided into control (-dBBr) and experimental (+dBBr) tubes. All subsequent steps were conducted in a dark environment to prevent photobleaching. Each tube received 5 µL of freshly prepared 0.2 M dibromobimane (dBBr, Sigma) or dimethyl sulfoxide (DMSO) and incubated for 1 hour at 4°C with gentle agitation. The crosslinking reaction was quenched with 50 µL of 0.2 M N-ethylmaleimide (NEM) for 30 minutes.

ASBT purification: Membrane proteins were solubilized in 0.5 mL membrane solubilization buffer (20 mM HEPES, pH 7.5, 150 mM NaCl, 10% glycerol, 2% DDM, 20 mM imidazole, 1 mM PMSF) through agitation (1000 RPM, 20°C) for 5 minutes, followed by a 15 minutes incubation on ice. The mixture was then centrifuged at 20,000*g* for 5 minutes at 4°C. The supernatant (800 µL) was incubated with 30 µL of Talon beads (Takara Bio) at 4°C for 1 hour with continuous shaking. The beads were washed twice with wash buffer (20 mM HEPES pH 7.5, 150 mM NaCl, 10% glycerol, 0.2% DDM, 20 mM imidazole) and eluted with 60 µL of elution buffer (1.5x non-reducing sample buffer, supplemented with 450 mM imidazole).

The samples were subjected to SDS-PAGE gel electrophoresis and initially imaged using a gel documentation system with a 365 nm excitation wavelength (BioRad). Post-imaging, the gel was stained with Coomassie Instant Blue and dried. If needed, quantitative analysis was performed to ensure equal protein loading for a second round of gel electrophoresis.

### Materials

FtsH antibody was obtained from National BioResource Project (NBRP)-E. coli, Japan Anti HA was from Sigma-Aldrich, H9658

Lysozyme was from Angene

Turbo DNase was from Jena biosciences cOmplete Protease Inhibitor was from Roche [^35^S]Methionine was from PerkinElmer

## Acknowledgments

We thank Daniel O Daley, University of Stockholm, Yoshinori Akiyama, Kyoto University and Ming Zhou, Baylor College of Medicine, for kindly providing plasmids. We thank Vasiliy Vladimirov for fruitful discussions, Yael Fridmann Sirkis for help with thermal unfolding, and Maja Rennig and Morten HH Nørholm for help with characterization of ASBT degradation. We are grateful to Gunnar von Heijne for initial financial support (grants by the Knut and Alice Wallenberg Foundation (2017.0323) and the Swedish Research Council (621-2014-3713) to GvH) and discussion, and for critically reading the manuscript. This research was generously supported by research grants from the ISRAEL SCIENCE FOUNDATION (grants no. 2207/21 and 2208/21), and the Center for New Scientists at the Weizmann Institute of Science. Illustrations were Created with BioRender.com.

## Author Contributions

M.C-D., H.P-Z and N.F. designed the study and the experiments.

M.C.D., N.R., T.O., M.P. and A.B. performed the experiments.

N.F., M.C-D., N.R. and T.O. conducted the majority of data analysis.

A.B.D.B performed Alphafold predictions.

N.F. and M.C.D. wrote the manuscript.

All authors discussed the results and contributed to editing the manuscript.

N.F. conceived the study and supervised the work.

---

**Fig. S1.**
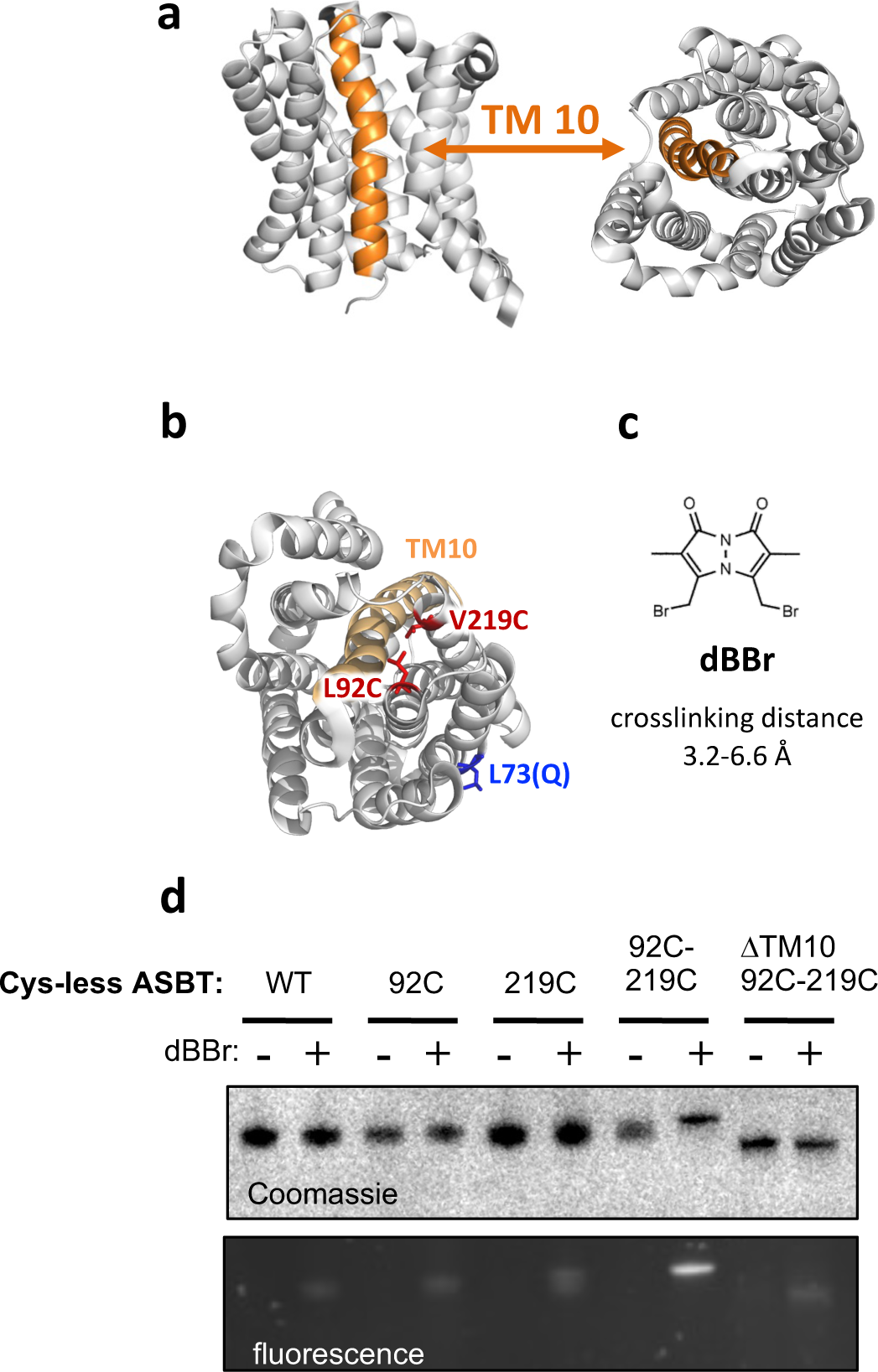
Probing the folding status of ASBT by Crosslinking. a. Crystal structure of ASBT (PDB: 4N7X), viewed from within (left) or above(right) the plane of the membrane. The central location of TM10 (orange) suggests that its deletion will likely cause misfolding. b. Location of the two engineered Cys in the ASBT structure. The crosslinked residues at positions 92 and 219 are located at the edges of TMs 4 and 8, which form extensive interactions with TM3, where L73Q lies, and TM10. Their proximity, as assayed by crosslinking, can serve as a measure for the proper folding of this structural module. c. Structure of the crosslinker dBBr, having a crosslinking distance of 3.2-6.6 Å. d. Crosslinking of ASBT by dBBr in native membranes confirms that two engineered cysteines are required for crosslinking. Crosslinking was demonstrated by a shift SDS-PAGE migration (Coomassie) and by the appearance of a fluorescent band of crosslinked protein upon the reaction of a single dBBr with two cysteines simultaneously. The indicated ASBT variants were cross-linked in native membranes, followed by purification and SDS-PAGE.

**Fig. S2.**
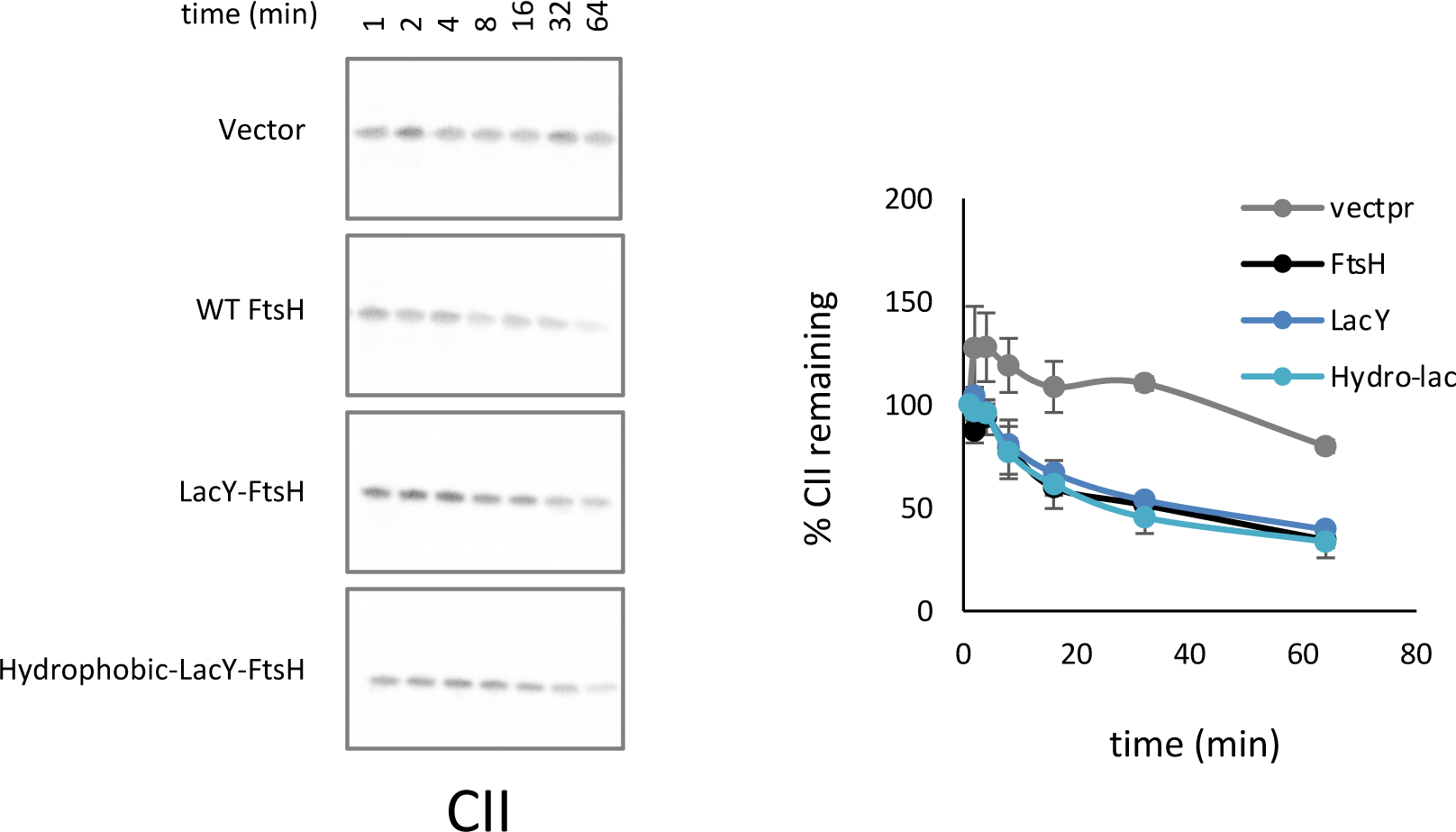
CII degradation by FtsH variants suggests that LacY-FtsH are active towards a soluble substrate. CII degradation was followed by radioactive pulse-chase followed by autoradiography. Left: representative gels showing the degradation of CII over time by different FtsH variants. Right: Quantifications of three experiments. Means ± SEM are shown.

**Fig. S2.**
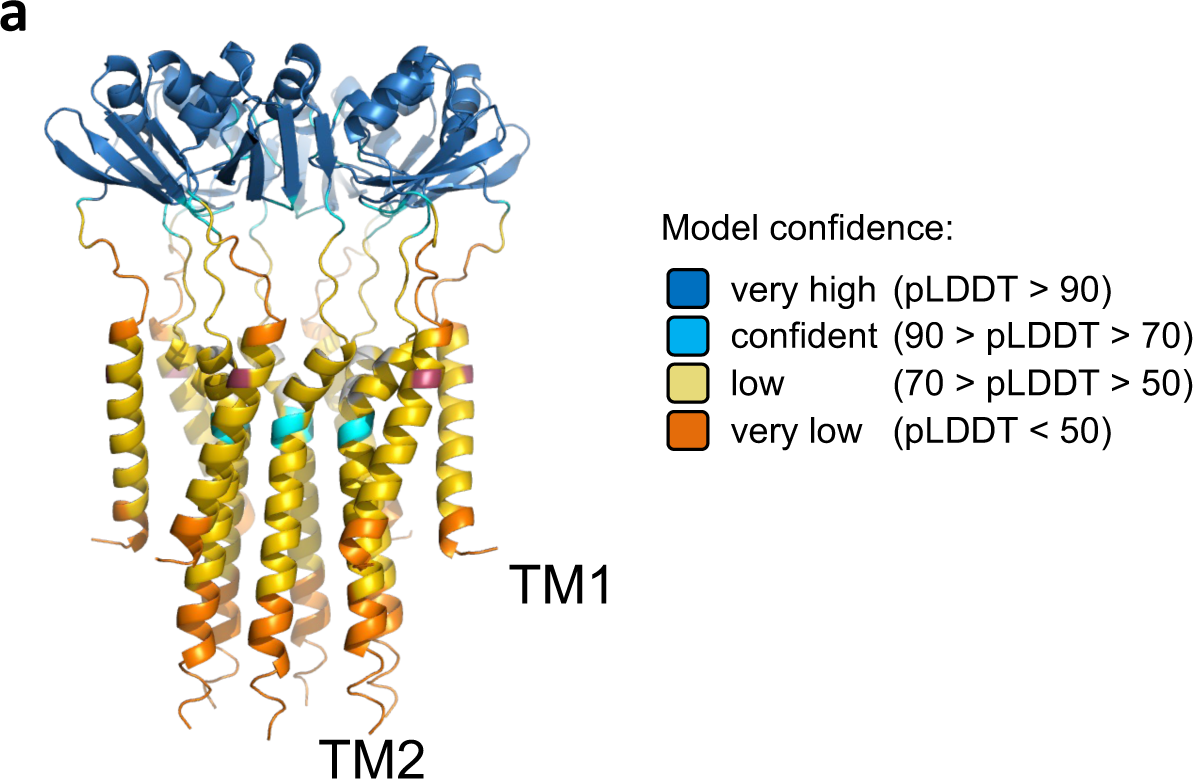
AlphaFold model of the TM and periplasmic domains of FtsH. TM1 is surrounding a ring of TM2 core helices. The model was generated using AlphaFold Multimer, using residues 1-137 of FtsH. Confidence scoring according to AlphaFold, suggests that the modeling of the periplasmic domain is more accurate than the TM domain.

**Fig. S4.**
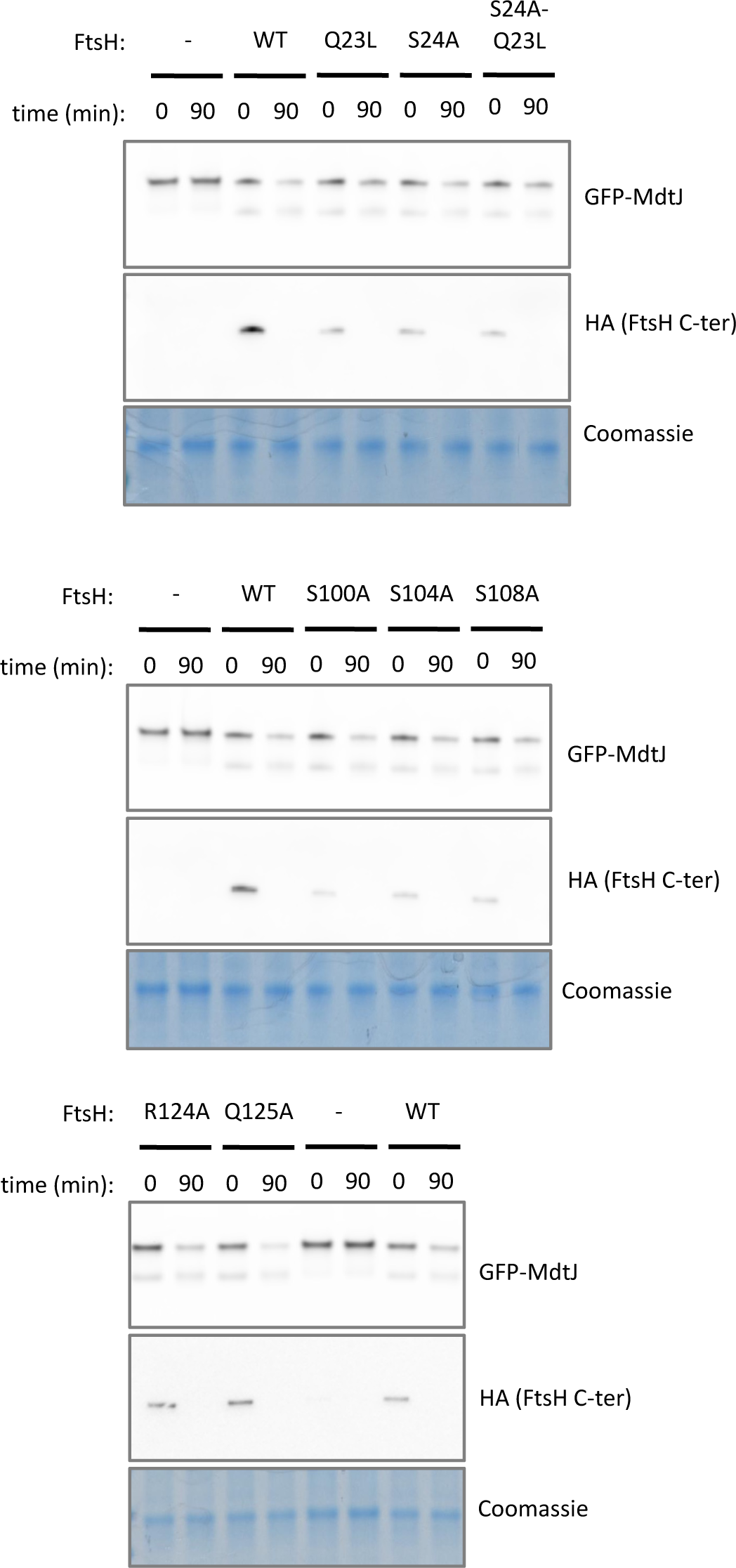
Representative SDS-PAGE gels showing the activity of FtsH variants in degradation of MdtJ and of the FtsH C-terminal HA tag. The degradation experiment was done similarly to Fig. 1a, with additional detection of FtsH C-terminus by anti-HA antibodies. Coomassie is shown as a loading control.

